# Foveal pRF properties in the visual cortex depend on the extent of stimulated visual field

**DOI:** 10.1101/2020.01.28.923045

**Authors:** Gokulraj Prabhakaran, Joana Carvalho, Azzurra Invernizzi, Martin Kanowski, Remco J. Renken, Frans W. Cornelissen, Michael B. Hoffmann

**Author notes:** Correspondent details: Michael B. Hoffmann, Universitäts-Augenklinik, Visual Processing Laboratory, Leipziger Str. 44, 39120 Magdeburg, Germany.

## Abstract

Previous studies demonstrated that alterations in functional MRI derived receptive field (pRF) properties in cortical projection zones of retinal lesions can erroneously be mistaken for cortical large-scale reorganization in response to visual system pathologies. We tested, whether such confounds are also evident in the normal cortical projection zone of the fovea for simulated peripheral visual field defects. We applied fMRI-based visual field mapping of the central visual field at 3 Tesla in eight controls to compare the pRF properties of the central visual field of a reference condition (stimulus radius: 14°) and two conditions with simulated peripheral visual field defect, i.e., with a peripheral gray mask, stimulating only the central 7° or 4° radius. We quantified, for the cortical representation of the actually stimulated visual field, the changes in the position and size of the pRFs associated with reduced peripheral stimulation using conventional and advanced pRF modeling. We found foveal pRF-positions (≤3°) to be significantly shifted towards the periphery (p<0.05, corrected). These pRF-shifts were largest for the 4° condition [visual area (mean eccentricity shift): V1 (0.9°), V2 (0.9°), V3 (1.0°)], but also evident for the 7° condition [V1 (0.5°), V2 (0.5°), V3 (0.9°)]. Further, an overall enlargement of pRF-sizes was observed. These findings indicate the dependence of foveal pRF parameters on the spatial extent of the stimulated visual field. Consequently, our results imply that, previously reported similar findings in patients with actual peripheral scotomas need to be interpreted with caution and indicate the need for adequate control conditions in investigations of visual cortex reorganization.

**Highlights:**

1. Foveal pRF properties change in controls with restricted peripheral stimulation
2. pRFs shift in position and enlarge in size for reduced stimulation extent
3. Alterations in pRF characteristics in patients should be interpreted with caution
4. Adequate control conditions needed in investigations of visual cortex plasticity

## 1. Introduction

Receptive field (RF) characteristics of neurons driven by visual input and their dynamics have for long been of fundamental interest in order to understand the mechanisms underlying visual processing. In contrast to invasive single-neuron electrophysiological recordings, functional MRI (fMRI) based RF measures reflect the aggregate characteristics of a population of neurons in a single voxel, termed population receptive field (pRF), where a pRF refers to the region in the visual field (VF) that elicits a response in the voxel. Over the last decade, a model based fMRI approach termed population receptive field mapping (Dumoulin and Wandell, 2008), has been widely used in investigating visual cortex functioning and contributed to our understanding of pRF characteristics of the visual cortex in healthy vision (Harvey and Dumoulin, 2011; Lee et al., 2013; Wandell and Winawer, 2015; Zeidman et al., 2018; Zuiderbaan et al., 2012). Furthermore, pRF modeling allowed us to quantitatively assess potential alterations of pRF characteristics in the visual cortex in the face of retinal lesions (Barton and Brewer, 2015; Baseler et al., 2011), developmental disorders (Ahmadi et al., 2019; Carvalho et al., 2019; Hoffmann et al., 2012; Hoffmann and Dumoulin, 2015) and trauma (Haak et al., 2014; Halbertsma et al., 2019; Papanikolaou et al., 2014). While alterations could be interpreted as evidence for potential cortical remapping and as an explanation for changes in fMRI responses following visual field defects (Baker et al., 2008; Dilks et al., 2009; Ferreira et al., 2017; Zhou et al., 2017), there is growing evidence for more conservative views on the nature and extent of adult visual cortex plasticity (Masuda et al., 2010, 2008; Wandell and Smirnakis, 2009). E.g., larger pRFs in the lesion projection zones (LPZ) in the primary visual cortex in patients with macular degeneration were also evident in controls with simulated lesions (Barton and Brewer, 2015; Baseler et al., 2011; Haak et al., 2012), thus questioning the concept of large-scale long-term reorganization in the visual cortex. While this bias has been well investigated in the lesion projection zone for deprived foveal stimulation (Haak et al., 2012; Morland, 2015), studies on the pRF dynamics in the normal projection zone associated of the fovea during deprived peripheral stimulation are currently emerging. In fact, recent investigations of patients with peripheral visual field defects, due to retinitis pigmentosa (RP) (Ferreira et al., 2017) or glaucoma (Zhou et al., 2017), report a displacement of pRFs in the normal projection zone, i.e., foveal pRFs to parafoveal or eccentric position. These shifts were taken as evidence for cortical remapping, in comparison to healthy controls with intact VF representations. While the authors of these studies acknowledge the limitations of a lack of appropriate control comparisons, it is unresolved whether the observed effects also occur for simulated peripheral visual field defects. This differentiation, however, is instrumental to dissociate long-term pathology-induced plasticity from short-term adaptation effects associated with the extent of visual stimulation.

In the present study, we intended to bridge this gap by reporting pRF measurements in healthy participants for three different stimulus sizes; a normal retinal representation of the visual space (14°) and two restricted representations comprising only the central 7° and 4° respectively. The size of the stimulus was reduced by applying two differently sized mean luminance masks in the peripheral visual field (> 7° and > 4°) of the 14° stimulus space. We investigated the effect of reduced peripheral stimulation on the pRF estimates of the central visual field representation in primary (V1) and extra-striate (V2 & V3) visual cortex. Our study revealed a pRF displacement towards the stimulus border and an enlargement of foveal pRFs (< 3°) for the restricted stimulus condition (7° and 4°), in comparison to the 14° pRF estimates. As possible explanations of these effects, we discuss the contribution of both physiological mechanisms and potential methodological and modeling causes.

## 2. Methods

### 2.1 Participants

Eight individuals (age: 21 to 28; 4 males and 4 females) with normal vision (best-corrected decimal visual acuity ≥ 1.0 (Bach, 1996)) took part in the study. All participants gave their informed written consent. The study was approved by the ethics committee of the University of Magdeburg and the procedure adhered to the tenets of the Declaration of Helsinki.

### 2.2 MR stimulus

The visual stimulus, programmed in MATLAB (Mathworks, Natick, Massachusetts, USA) using Psychtoolbox-3 (Brainard, 1997; Pelli, 1997), was projected (resolution: 1920 × 1080 pixels) to a screen at the rear end of the magnet bore. Participants viewed the stimulus monocular with their dominant eye at a distance of 35 cm through an angled mirror. The stimulus was a checkerboard pattern (mean luminance: 109 cd/m^2^; contrast: 99%; check size: 1.57°) exposed through a bar aperture (3.45°) moving in eight different directions (2 horizontal, 2 vertical and 4 oblique orthogonal) presented over four trials within a scan. The temporal sequence of each trial comprised of a sweep in one of the horizontal or vertical direction (24s) and a sweep in an orthogonal direction (12s) followed by mean luminance gray (12s). The bar moved at a step rate of 2°/s across a circular aperture subtending 14° radius (the maximum possible stimulus size with our current fMRI setup). In addition to the 14° condition, we estimated in separate scans during the same session the pRFs for two other stimulus size conditions, i.e., 7° and 4° radius, by masking the peripheral section of the 14° stimulus (Fig 1). Hence we refer the 7° and 4° stimulus conditions as masked conditions and the 14° stimulus as the unmasked condition. The spatial and temporal properties of the stimulus remained same for all the three conditions but only the extent of stimulation varied. The duration of each of these conditions was 192 seconds. Each condition was repeated four times in an interleaved design. The participants were instructed to focus their attention on a small fixation cross placed at the center of the stimuli and report a change in the fixation color via button press.

**Fig 1:**
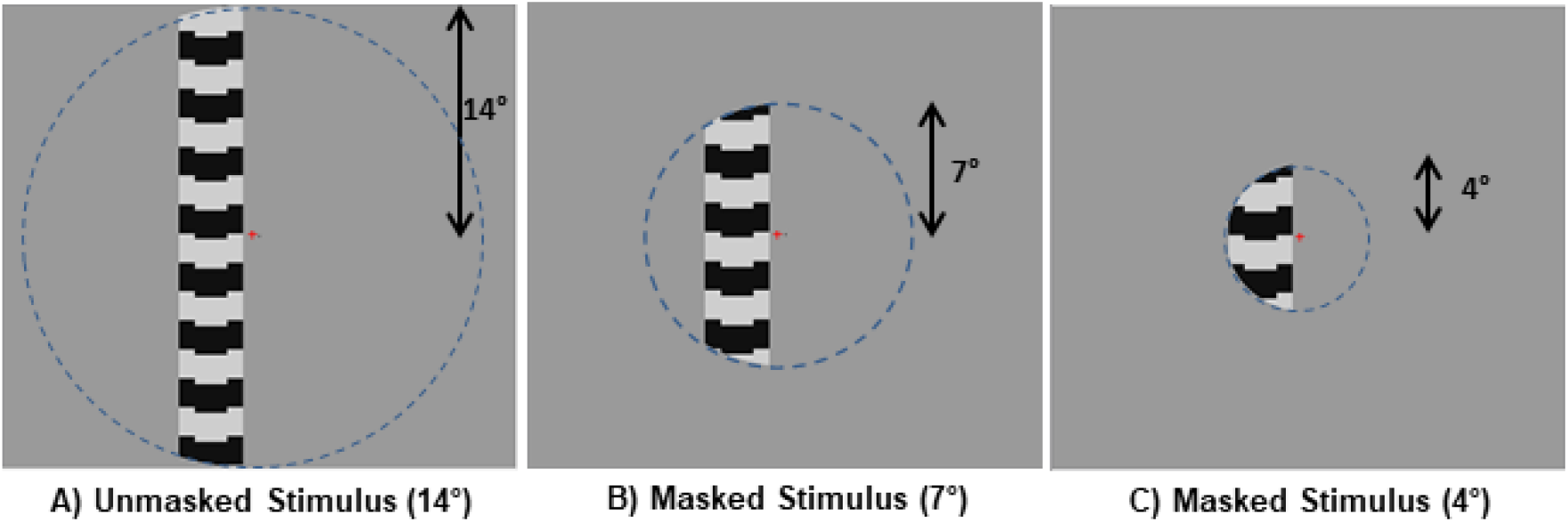
Illustration of the three stimulus size configurations; unmasked (**A**) and the masked stimulus (**B** & **C**) configurations. The size of the unmasked stimuli was 14° (radius) and was restricted for the masked stimuli by simulating mean luminance masks in the peripheral VF (> 7° and > 4°) of the 14° stimulus space, stimulating only the central 7° and 4° respectively. Blue dashed boundaries indicate the spatial extent of the stimulated visual field (depicted here only for illustration purpose and was not visible during stimulus presentation) during the different conditions, whereas the spatial and temporal properties of the stimuli did not change. Participants fixated their attention on a small cross placed at the center of the stimuli and reported a change in the fixation color using a button press.

### 2.3 MRI acquisition

All MRI measurements were obtained with a 3 Tesla Siemens Prisma scanner using only the lower part of a 64-channel head coil. This results in a 34-channel coil covering most of the brain while allowing an unrestricted view to the projection screen. fMRI scans were acquired using a T2*-weighted BOLD gradient-EPI sequence (TR | TE = 1500 | 30 ms & voxel size = 2.5^3^ mm^3^). The first 8 volumes from each scan were removed automatically by the scanner to allow for steady magnetization. Each scan comprised a total of 136 volumes. One anatomical T1-weighted scan (MPRAGE, 1 mm isotropic voxels, TR | TI | TE = 2500 | 1100 ms | 2.82 ms) was collected for each participant to allow for gray-white matter segmentation and surface based analyses.

### 2.4 Data preprocessing and analysis

T1-weighted anatomical scans were segmented for gray-white matter boundaries using Freesurfer (https://surfer.nmr.mgh.harvard.edu/) and manually corrected for possible segmentation errors using ITK gray software (https://github.com/vistalab/itkgray). The gray-white boundaries were reconstructed to generate a 3-dimensional rendering of the cortical surface (Wandell et al 2000). fMRI scans of each individual participant were corrected for within and between scan head motion artefacts using AFNI (https://afni.nimh.nih.gov/). For each participant, motion-corrected fMRI time series of the different stimulus size conditions (14°, 7° and 4°) were averaged together into separate groups with MATLAB based Vistasoft tools (https://github.com/vistalab/vistasoft) and were aligned spatially to the anatomical scan using Kendrick Kay’s alignment toolbox (https://github.com/kendrickkay/alignvolumedata).

### 2.4 pRF modeling

We estimated the pRF parameters independently for our three stimulus conditions using a custom implemented advanced pRF modeling method based on Bayesian inference and Markov Chain Monte Carlo (MCMC) sampling (Adaszewski et al., 2018; Carvalho et al., 2019; Zeidman et al., 2018). In a comparative approach, we also looked at the conventional pRF estimates (Dumoulin and Wandell, 2008) to inspect for any model specific variations in the estimates. Since the Bayesian pRF modeling builds on the conventional method, we start with the description of the latter’s procedure.

#### 2.4.1 Conventional pRF

For each voxel, the voxel’s fMRI time-series was used to estimate the aggregate receptive field properties of the underlying neuronal population using a 2D-Gaussian pRF model (described by three stimulus-referred parameters; position preferred in the visual field (x and y in Cartesian coordinates) and the spatial spread (σ)). The model, along with the time course of the stimulus, predicts the voxels fMRI response of a voxel and convolves with a canonical hemodynamic response function (HRF) (Friston et al., 1998). Approximately 100,000 plausible combinations of pRF parameters (x, y,σ) were generated to compute predictions of the observed BOLD time-series of each voxel. The optimal pRF parameters, best fitting the predicted and actual voxel time-series were estimated by minimizing the sum of squared errors (RSS) between the two. Position parameters were used to derive pRF 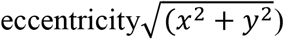 and polar angle 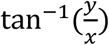 and pRF size from the spatial spread of the fitted 2D Gaussian model.

#### 2.4.2 Bayesian MCMC

As we followed the nomenclature and mathematical notations for the Bayesian modeling (Zeidman et al., 2018) and MCMC sampling (Adaszewski et al., 2018; Carvalho et al., 2019) used elsewhere, we present here a more generic description of the method. With the conventional pRF method, we have fixed model parameters and use the variation between the predicted and the observed data to infer the most probable estimates of a voxel without any information on the probability of the estimates. In contrast to this, the Bayesian approach computes the posterior probability (*posterior*) of the predicted pRF parameter combination. The posterior follows the Bayes’ rule (Equation 1) translating as the probability of a pRF parameter combination (location: x, y and spread σ) given the observed BOLD time series (BOLD_TS_), hemodynamic response function (HRF) and the stimulus representation (Π).

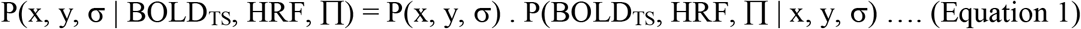

We computed the posterior as the product of probability of a parameter combination (prior) and the probability of the observed voxel response given a parameter combination (likelihood). The likelihood of a parameter combination was computed as the probability density function of the error between the predicted and the observed voxel time-series given a set of pRF parameters (as in conventional pRF). We restricted the plausible pRF position (x, y) parameter to the area of our unmasked stimulus (14° radius), whereas for the conventional pRF the field of view ranged between [−2*radius, 2*radius], i.e. [−28° to 28°], resulting in the pRFs to be placed beyond the stimulated visual field. To start with, we initialized the pRF position parameters with a non-flat prior which accounts for the cortical magnification factor i.e. we assume a higher probability for the pRFs to be centered foveally than in a peripheral position. However, for the pRF size (σ) parameter, a flat prior with equal probability for all possible widths within the permissible range [0.5° to 14°] was assigned.

We used the Markov Chain Monte Carlo (MCMC) approximation to efficiently sample the visual cortex, i.e. areas of the visual field with higher variance explained were more densely sampled. Per voxel 17500 samples were computed, which form the posterior distribution. We start with a set of initial pRF parameters (x, y, σ) and propose a new set of parameters based on the initial ones. The MCMC algorithm followed a coarse to fine approach for choosing the proposed parameters i.e. the step size was larger for the initial samples whereas it reduced with the iterations. We compute and compare the posteriors for the initial and the proposed parameters. If the proposed parameters adhere to the observed data better than the initial parameters, we accept the proposed parameters and assign the combination as the new initial parameters; else there is still a chance to accept the proposed parameters. For this a probability of random acceptance was defined, if the difference in likelihood between the proposed and initial parameters is bigger than the N(0,1), the parameters are updated. As we repeated this procedure for 17500 iterations, the parameters start to converge around the true measures underlying the neuronal population of a voxel. Our priors for the parameter combinations were not based on empirical data, which might result in the initial posterior samples to deviate from the true measure and skew the posterior distribution. To mitigate this possible bias, we discarded the posteriors of the initial 10% (1750) samples for further analysis. The Bayesian approach, for each voxel provides us a quantitative inference of the underlying distribution of the pRF model parameters (uncertainty). We used the full width at half maximum (FWHM) of the posterior distribution of the parameters as our measure of uncertainty, thereby investigating any changes in the behavior of the posterior distribution of each voxel for the different stimulus size conditions.

### 2.5 ROI definition and statistical analysis

Polar angle maps from the unmasked 14° condition for each participant were projected onto an inflated cortical surface. We delineated the borders of the primary (V1) and extra-striate (V2 and V3) visual cortex by following the phase reversals in the polar angle data (Sereno et al., 1995). All the further region of interest (ROI) analyses and statistics were performed with custom written scripts and statistical toolbox functions in MATLAB. Only voxels with an explained variance above 15% for the unmasked stimulus condition (14°) were included for all the subsequent analysis presented here (applying no threshold did not influence the findings we report here).

## 3. Results

### 3.1 Cortical representation of the stimulus

Firstly, we examined the representation of the Bayesian derived pRF eccentricities (preferred position in the visual field VF) in the cortex for the different stimulus conditions. For the unmasked 14° stimulus, the maps from all the participants followed a retinotopic organization in the primary (V1) and extra striate visual cortex (V2 and V3) spanning the entire stimulus radius of 14°. Whereas with masked 4° and 7° conditions, as expected, we observed a restricted representation in the cortex, in congruence with the stimulated VF, i.e. spanning a reduced eccentricity range. This is depicted for a representative participant in Figure 2A. In addition to the restriction of the representation, we also observed a trend in the voxels’ pRF centers to be more eccentric for the 4° and 7° conditions than for the 14° stimulus. This indicates an attraction of these pRFs towards the stimulus border. We illustrate this change in a sample participant by defining a small random paracentral ROI in V1 depicted by the blue borders in Fig 2A. The ROI included only those voxels which were stimulated for all three stimulus conditions. Fig 2B shows the plot of the mean preferred eccentricity of the voxels in the ROI for the three stimulus sizes. For the 14° stimulus condition, the mean preferred eccentricity in the VF was 1.5°, whereas for the 7° and 4° conditions, it increased to 2.1° and 3.1° respectively. The plots clearly show that for the masked conditions pRF positions were more eccentric than for the 14° condition. This effect was larger for the smallest stimulus size, i.e. for the 4° condition.

**Fig 2:**
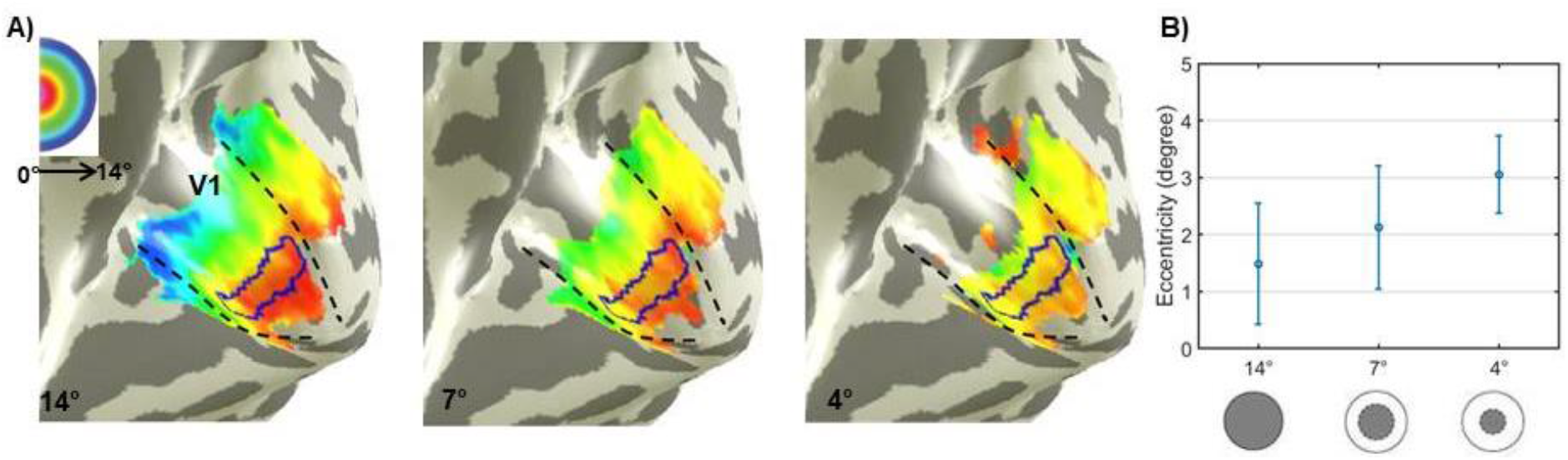
Bayesian MCMC derived pRF eccentricities **(A**) of a representative participant mapped on inflated right visual cortex for the different stimulus sizes - 14° (unmasked), 7° and 4° (masked). Dashed black lines delineate the primary visual cortex V1. False-color representation from dark orange to dark blue in the unmasked stimulus illustrates the retinotopically organized eccentricities from 0° to 14°. Restricted cortical representation and changes in the pseudo-color progression are visualized in the masked condition maps, respective to their stimulus size. Panel **B** plots the mean and standard deviation of eccentricities within a small randomly defined paracentral ROI in V1 (indicated by the blue boundaries in the map) for the three stimulus sizes. Insets at the bottom of panel B shows a sample depiction of the stimulus configurations (grayed section represents the stimulated visual field). The plots clearly demonstrate the increase in eccentricity upon the restriction of the stimulus size.

### 3.2 Stimulus size dependent changes in pRF properties

#### 3.2.1 Displacement of pRF preferred position

Next we assessed the spatial extent of these observed shifts in the pRF position with reduced stimulus size at the group level. For this purpose, we grouped the voxels into bins of 1° based on the 14° Bayesian derived pRF eccentricities, and determined for each visual area (V1, V2, and V3) and bin, the mean shift in eccentricity for the 4° or 7° conditions in comparison to the unmasked condition (Eccentricity_14°_ – Eccentricity_4° or 7°_) (Fig 3; mean eccentricity shift (n=8) plotted with standard error of the mean (SEM)). Due to our interest in understanding the pRF dynamics in the stimulated VF, we restricted our further analysis to the stimulated eccentricities of the restricted stimulus conditions (i.e. 4 bins for the 4° and 7 bins for the 7° conditions). For the 4° condition, we observed the mean eccentricity of the bins to be more eccentric than for the 14° condition and the difference was, as expected, negative for the stimulated eccentricities (Fig 3A). This increase was particularly pronounced for the central eccentricities (< 3°) in contrast to those adjacent to the border of the stimulus, i.e. 3° to 4°. The trend progresses similarly across V1, V2 & V3. Although we report a similar position displacement for the 7° condition (Fig 3B) for the 1° and 2° bins, the shifts were smaller than the 4° condition and did not propagate across the further eccentricities. A similar pattern of eccentricity shifts was also observed for the estimates from the conventional pRF approach (See Supplementary Fig S1 A&B). Subsequently, we performed independent 2-way ANOVAs [Factor 1 *eccentricity* (bins), Factor 2 *model* (Bayesian pRF and conventional pRF)] on these reported shifts for each visual area (V1, V2 & V3) and stimulus combinations (14°-4° and 14°-7°; Table 1). The ANOVAs revealed a main effect of *eccentricity* in the reported shifts (Table 1A). Although, the shifts observed for the 7° stimulus condition were larger for the Bayesian approach than for conventional pRF-mapping, the ANOVAs did neither show a significant effect of *model* to the displacements (p > 0.093 (with an exception of marginal significance for 1/6 ANOVAs; p>0.04)) nor an interaction of *eccentricity* and *model* (p > 0.75) in any of the visual areas (see Table 1A for region and stimulus condition specific statistics). Our post-hoc 2-sample t-tests with Holm-Bonferroni correction (Holm, 1979) showed the observed shifts with 4° stimulus size to be significant (p < 0.05) for the eccentricities < 3° for all visual areas, except for V3 with significant shifts only for eccentricities < 2°. For the 7° stimulus condition, pRFs were significantly displaced only for the central 2 degrees.

**Fig 3:**
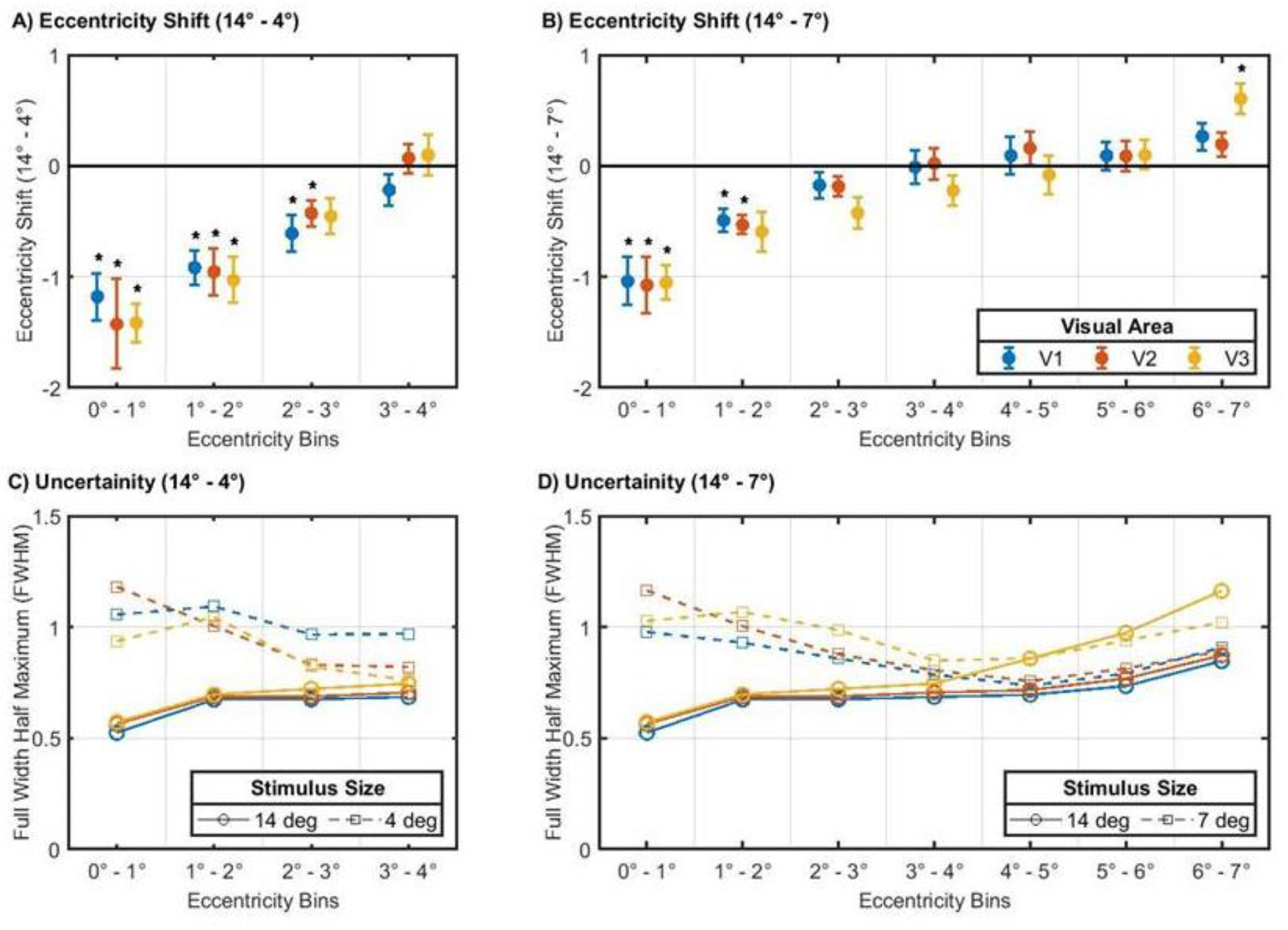
Differences in Bayesian-MCMC-derived pRF-eccentricity measures, i.e. unmasked 14° condition minus the masked smaller stimulus conditions, i.e. (**A)** Eccentricity_(14° − 4°)_ **(B)** Eccentricity_(14° − 7°)_. For each participant, voxels were grouped into bins of 1° based on their 14° condition eccentricities (x-axis). All the analysis was restricted only to the stimulated eccentricities of the restricted stimulus conditions (i.e. 4 bins for the 4° and 7 bins for the 7° conditions) and included only those voxels which explained at least 15% of the time-series variance for the unmasked configuration. In the y-axis, for each eccentricity bin, the group level (n=8) mean shift in eccentricity and standard error are plotted for visual areas, V1 (blue), V2 (Red) and V3 (orange). Shift of the foveal eccentricities (<3°) in the negative direction implies a more eccentric preferred pRF position for the 4° or 7° stimulus compared to the 14° condition. The shifts were larger for the 4° stimulus than the 7° stimulus and similar across V1, V2 and V3. Visual areas, eccentricity bins and stimulus conditions with significant effects (p<0.05) after Holm-Bonferroni correction are indicated by “*”. Bottom panels (**C&D**) shows the bin-wise comparison of FWHM of the eccentricity distribution (Bayesian measure of uncertainty) for the 14° (circle) vs. 4° or 7° (square) conditions. Note the increased uncertainty for the masked conditions, in particular for the bins with larger shifts.

**Table 1:**
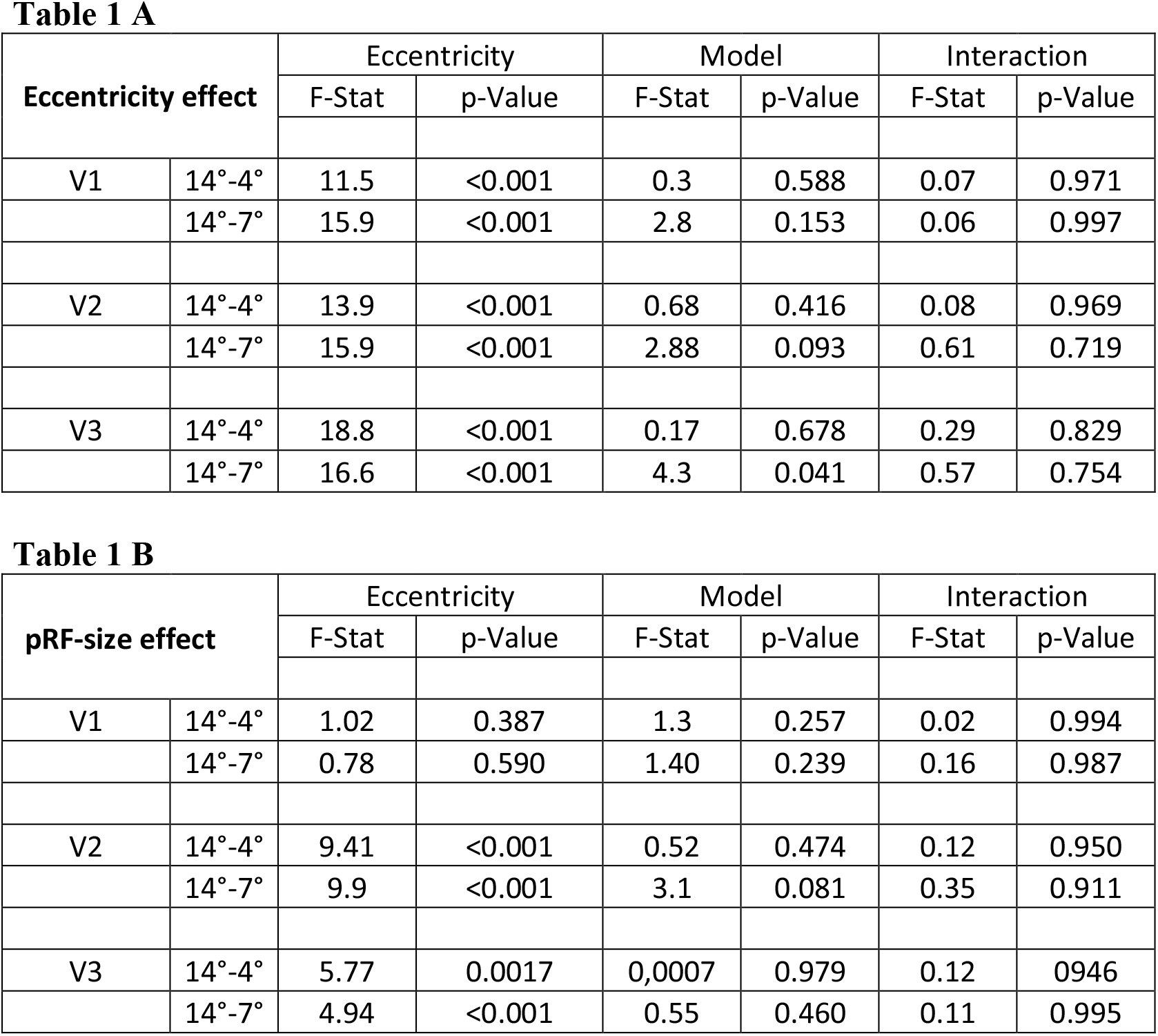
Two-way ANOVA of the reported eccentricity (A) and pRF size (B) shifts for each visual area (V1, V2 & V3) and stimulus combinations (14°-4° & 14°-7°). Factor 1: eccentricity (bins); Factor 2: model (Bayesian pRF or conventional pRF). Only the bins within stimulated eccentricities were used in the analysis.

| Eccentricity effect |  | Eccentricity |  | Model |  | Interaction |  |
| --- | --- | --- | --- | --- | --- | --- | --- |
|  |  | F-Stat | p-Value | F-Stat | p-Value | F-Stat | p-Value |
| V1 | $14^\circ$-$4^\circ$ | 11.5 | <0.001 | 0.3 | 0.588 | 0.07 | 0.971 |
| | $14^\circ$-$7^\circ$ | 15.9 | <0.001 | 2.8 | 0.153 | 0.06 | 0.997 |
| V2 | $14^\circ$-$4^\circ$ | 13.9 | <0.001 | 0.68 | 0.416 | 0.08 | 0.969 |
| | $14^\circ$-$7^\circ$ | 15.9 | <0.001 | 2.88 | 0.093 | 0.61 | 0.719 |
| V3 | $14^\circ$-$4^\circ$ | 18.8 | <0.001 | 0.17 | 0.678 | 0.29 | 0.829 |
| | $14^\circ$-$7^\circ$ | 16.6 | <0.001 | 4.3 | 0.041 | 0.57 | 0.754 |

| pRF-size effect |  | Eccentricity |  | Model |  | Interaction |  |
| --- | --- | --- | --- | --- | --- | --- | --- |
|  |  | F-Stat | p-Value | F-Stat | p-Value | F-Stat | p-Value |
| V1 | $14^\circ$-$4^\circ$ | 1.02 | 0.387 | 1.3 | 0.257 | 0.02 | 0.994 |
| | $14^\circ$-$7^\circ$ | 0.78 | 0.590 | 1.40 | 0.239 | 0.16 | 0.987 |
| V2 | $14^\circ$-$4^\circ$ | 9.41 | <0.001 | 0.52 | 0.474 | 0.12 | 0.950 |
| | $14^\circ$-$7^\circ$ | 9.9 | <0.001 | 3.1 | 0.081 | 0.35 | 0.911 |
| V3 | $14^\circ$-$4^\circ$ | 5.77 | 0.0017 | 0.0007 | 0.979 | 0.12 | 0.946 |
| | $14^\circ$-$7^\circ$ | 4.94 | <0.001 | 0.55 | 0.460 | 0.11 | 0.995 |

In addition to the eccentricity measures, we compared the distribution of Bayesian derived eccentricities for each bin for the 14° condition with either the 4° or 7° condition (Fig 3C (14°-4°) & 3D (14°-7°)). As detailed in Methods, we used the FWHM of the distribution as our measure of uncertainty, as this reflects the range of RF positions represented by the neuronal population within each bin. As expected, the FWHM increased for the masked conditions over the unmasked condition especially in the central eccentricities. The wider distribution in the center with reduced stimulus size indicates that the neuronal populations in these voxels now represent a larger range of VF positions and the estimates obtained for these conditions are most likely to be biased. There is a high correspondence of larger uncertainty for the eccentricity bins with the biggest displacement.

As our measures show the aggregate estimates of all the voxels in a bin, the shifts observed could be the result of either (1) few voxels having very large displacements or (2) a substantial number of voxels with moderate displacements. To deduce the percentage of voxels contributing to the reported shifts, we defined a ROI with voxels representing the central eccentricities (<3°) in the 14° stimulus size condition and performed a voxel-wise comparison of the preferred pRF eccentricity between the three different conditions. Based on this comparison, we classified the voxels in the ROI into three categories; (1) voxels with at least 1/3rd increase in the pRF eccentricity measure for the 4° or 7° stimulus in comparison to the 14° condition *Eccentricity*_*change* ≥ *33*%_), (2) voxels with at least 1/3rd decrease in the pRF eccentricity measure (*Eccentricity*_*change* ≤ *33*%_), and (3) the remaining voxels as ones with no substantial position change between the conditions *Eccentricity*_*no-change*_. We observed nil or a very negligible number of voxels with Eccentricity_*change* ≤ *33*%_), for the masked conditions compared to the 14° condition. Fig 4 shows the percentage of voxels with *Eccentricity_change ≥ 33%_* (mean of all the participants (n=8) and the SEM). For the central representation, we observed approximately 50% of voxels contributing to the position displacement for the 4° masked conditions and 35% for the 7° condition across the visual hierarchy, indicating that these shifts are not driven by a very few voxels with ectopic RFs.

**Fig 4:**
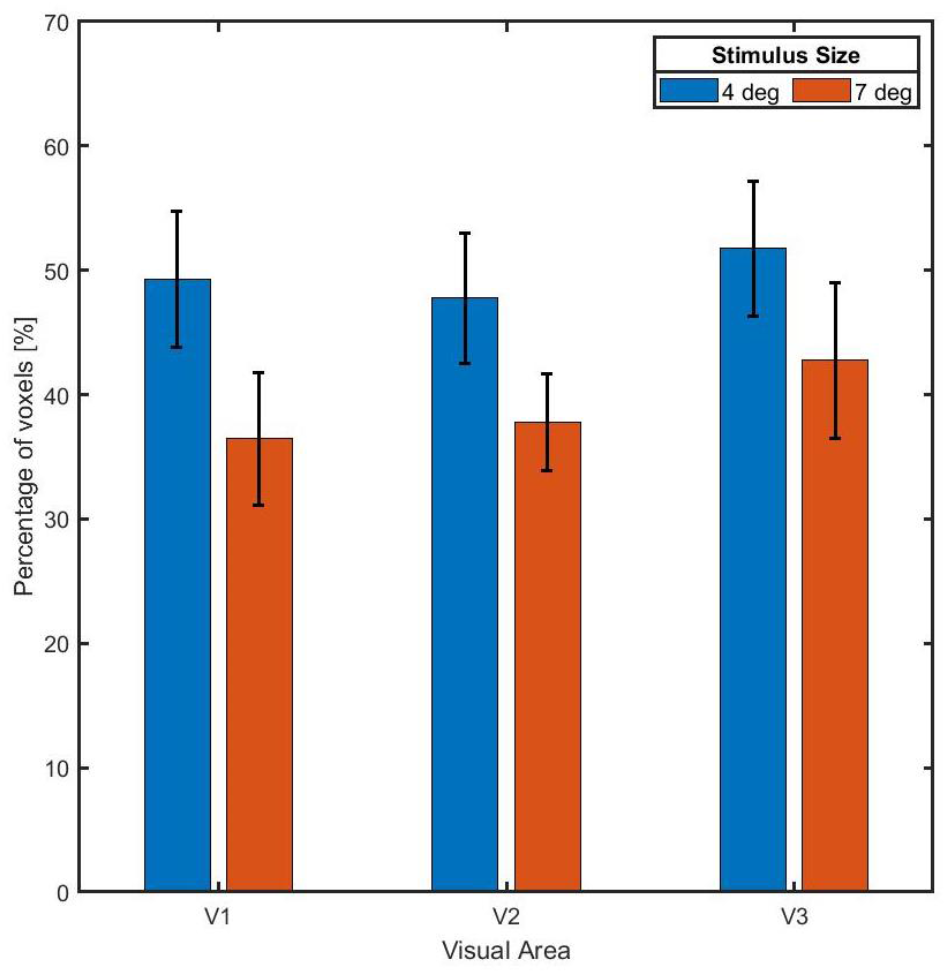
Percentage of voxels contributing to the observed foveal pRF displacements. Bayesian derived pRF eccentricities of voxels representing the foveal eccentricities (< 3°) for the unmasked 14° condition were compared voxel-wise with their smaller stimulus condition estimates (4° or 7°). Plot shows the mean percentage of voxels (n=8) with at least a 1/3^rd^ increase in the pRF eccentricity measure (Eccentricity_change ≥ 33%_) for the 4° (blue bars) and 7° (red bars) stimulus in comparison to the 14° condition. Approximately 50% and 35 % of voxels contribute for the reported position displacements in the 4° and 7° configurations for V1, V2 and V3.

#### 3.2.2 Enlargement of pRF-size

After the analysis of the pRF position, we analyzed whether Bayesian derived pRF sizes are also affected by stimulus size. Fig 5A & 5B shows the bin-wise mean pRF-size shifts between the 14° and the 4° or 7° conditions. Similar to the preferred pRF position, we observed increased pRF-size estimates in the stimulated eccentricities for smallest stimulus size (4°), whereas it was very negligible for the 7° condition. The shifts decreased as a function of eccentricity and we did not observe any systematic pattern across the visual areas. Conventional pRF-size estimates also exhibited comparable shifts with reduced stimulus size (supplementary Fig S1 (C&D). We performed a 2-way ANOVA [Factor 1 *eccentricity* (bins), Factor 2 *model* (Bayesian pRF and conventional pRF))] (Table 1B). The effect of *model* did not reach significance. A main effect of *eccentricity* was evident for the observed size-shifts for V2 and V3, but not for V1 (for both 14°-4° and 14°-7°). For the 4° condition, post-hoc t-tests corrected for multiple comparison showed significant shifts in V2 (eccentricity bins 1 to 3) and V3 (bin 1), and for the 7° condition, the shifts in V2 (bins 1 to 5) and V3 (bins 1 to 3) were significant with the Bayesian pRF approach. Similar to pRF eccentricity, the uncertainty i.e. FWHM of pRF-size distribution was larger for the masked condition (Fig 5C & 5D).

**Fig 5:**
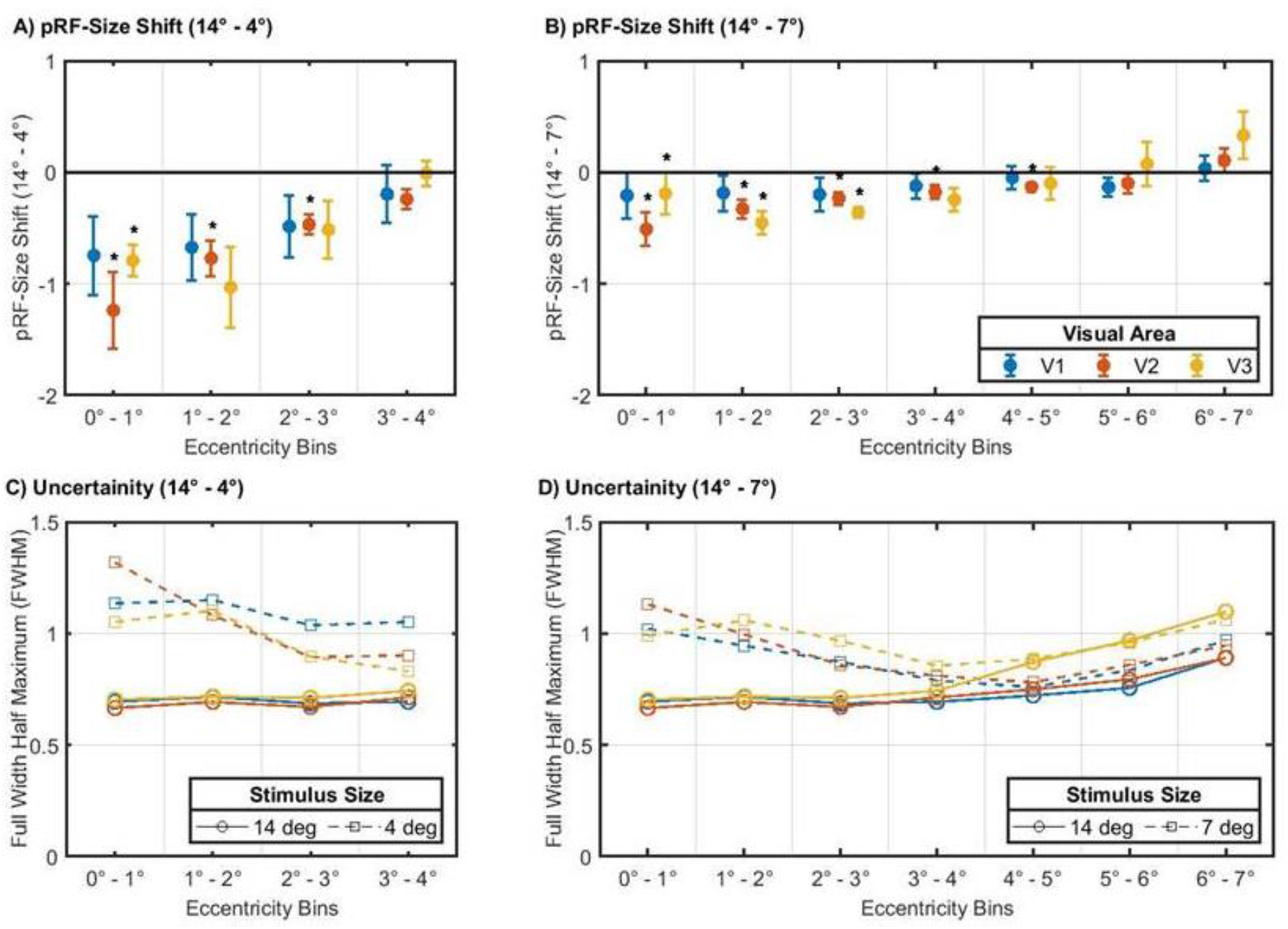
Differences in Bayesian-MCMC-derived pRF-size upon reduced stimulation extent. Differences calculated as (A) pRF-size(14° - 4°) (B) pRF-size(14° - 7°) for V1 (blue), V2 (Red) and V3 (orange). Grouping of voxels into eccentricity bins was performed as for Fig 3. For the y-axis the group level (n=8) mean shifts in pRF-size and standard error of the mean (SEM) are plotted. A change in the negative direction implies larger pRF sizes for the 4° or 7° stimulus compared to the 14° stimulus. We report larger absolute shifts for the 4° stimulus, whereas the 7° shifts were very small. Visual areas, eccentricity bins and stimulus conditions with significant effects (p<0.05) after Holm-Bonferroni correction are indicated by “*”. Bottom panels (C & D) show the bin-wise comparison of FWHM of the pRF-size distribution (Bayesian measure of uncertainty) for the 14° (circle) and 4° or 7° (square) conditions. Increased FWHM measures can be seen for the bins with larger pRF-size shifts, in particular for the 4° estimates.

#### 3.3 Modeling restricted stimulus representation

All the above reported analysis for all the three different stimulus sizes was modeled assuming an unmasked (14°) stimulus condition. As a sanity check, we assessed the presence of the observed effects while explicitly incorporating the actual stimulus representation in the models for the smaller stimulus size conditions. We did not observe any reduction of the stimulus size effects for this approach. Actually, we report shifts that were not restricted to the central eccentricities and that exceeded those found for our original approach, i.e. no modeling the reduced stimulus representation (see Supplementary Fig S2). Taken together, the observed effects were not reduced of knowledge about the simulated visual field defects entered the model.

## 4. Discussion

The findings from our study revealed, 1) shifts (displacement and enlargement) of the foveal pRFs (< 3°) from their preferred positions and size as estimated with 14° stimulus, when reducing the stimulus size, 2) shifts for the 4° exceeding than those for the 7° stimulus condition, 3) a propagation of the shifts across V1, V2 and V3, but without any hierarchical trend, 4) no significant differences between the modeling approaches (Bayesian pRF or conventional pRF). The novelty of our findings is that they were observed in the normal projection zone of the fovea, as opposed to reports on changes of the pRF-properties in the lesion projection zone of the damaged fovea (Dumoulin and Knapen, 2018; Morland, 2015; Wandell and Smirnakis, 2009). Similar changes in the receptive field estimates of the representation of the fovea in the early visual areas were reported by some studies on patients with peripheral visual field deficits, including RP (Ferreira et al., 2017) and glaucoma (Zhou et al., 2017). The authors proposed that the observed shifts are evidence of cortical reorganization in these patients, but at the same time in their interpretation acknowledged the possibility of control biases. Our results demonstrate that such alterations in the pRF estimates can also be observed in healthy individuals via mimicking scotomas of patients with peripheral retinal pathologies. This suggests that the previous findings in patients reflect normal cortical response behavior in a context of reduced peripheral stimulation. Consequently, our findings emphasize that care needs to be exerted before interpreting alterations in receptive field properties in patients with VF restrictions as definitive evidence for cortical reorganization. In particular, appropriate controls, e.g. with simulated visual field defects, are of great value investigating the scope of cortical plasticity.

A critical question prompted by the above findings concerns the nature and origin of the observed pRF estimate dependence on stimulus size. The aim of our study was to check the existence of such stimulus size dependent biases in pRF mapping. The next step is to identify the underlying causes and mechanism in future studies. Below, we suggest and discuss plausible explanations of the observed effects, i.e., (1) potential methodological or modeling biases associated with pRF mapping and (2) physiological mechanisms, i.e. changes in neuronal receptive field properties.

### 4.1 Modeling and stimulus configuration

Previous studies have reported biased pRF estimates in the neighborhood of the boundaries of simulated foveal (Baseler et al., 2011; Binda et al., 2013; Haak et al., 2012) and quadrantopic scotomas (Papanikolaou et al., 2015) in healthy individuals. Binda and colleagues (2013) proposed that such biases could be mitigated by taking into account the scotoma in the pRF model and using a randomized multifocal stimulus. Importantly, for the predominantly used non-randomized size-invariant bar stimulus, modeling the scotoma did not have a substantial effect on the biased estimates. This is consistent with our findings for the explicit inclusion of the reduced stimulus representation in the estimation models. We report stimulus size dependent pRF shifts that are not restricted to foveal eccentricities and larger than those observed when assuming an unmasked stimulus condition for modeling (Fig S2). Another study comparing different stimulus configurations for pRF mapping (Alvarez et al., 2015) illustrated eccentricity scaling and use of polar rather than Cartesian stimuli, to have a significant effect on the goodness of fits and pRF size estimates. Similar to our study, they also reported a display size bias in the pRF size by changing the viewing distance (but not by masking the stimulus periphery). Larger pRFs were observed in participants who experienced a 16° display (n=2) compared to those who experienced a 9° display (n=6). Although these results were in contrast to our finding of a mean expansion of the pRFs (at the group level) for the smaller stimulus size, we noticed heterogeneous single subject results (especially in V1) for the conventional pRF approach. Three out of our eight participants had smaller pRFs for the masked vs. unmasked stimulus condition (see SEM in Fig S1 (C&D), in particular in V1). However, this heterogeneity was not observed for the Bayesian model (see SEM in Fig 5 (A&B)), where all the participants showed pRF-size increase with reduced stimulus size, indicating the utility of the Bayesian approach in mitigating some methodological biases of conventional pRF modeling. Despite the possibility that biases associated with modeling and stimulus configurations could result in the shifts we report here, there are always tradeoffs in choosing the optimal stimulus design or modeling approach, e.g., the low power of multifocal stimulus (Binda et al., 2013; Ma et al., 2013), high predictive power but reduced accuracy of polar-coordinate based wedge and ring stimuli (Alvarez et al., 2015) or limitations of ascertaining absolute scotomas in patients to be included in the model. As over-manipulation of the stimulus and modeling characteristics can also limit the replicability and reproducibility of studies. The effective approach to circumvent potential methodological biases would be to match the experimental conditions between the controls and patients as closely as possible.

### 4.2 Ectopic receptive fields

As fMRI based pRF estimates reflect only the aggregate RF properties of all the neurons in the voxel, the observed shifts are possible even if the response characteristics of only a subset of the neuronal population changes. Haak and colleagues (2012) observed an activation of very few foveal voxels in healthy individuals even when only the peripheral VF was stimulated and highlighted the existence of neurons with ectopic receptive fields (RF) in the regions of cortex representing the central VF. Even though our masked stimulus might silence this ectopic neuronal subpopulation, it is highly unlikely that this mechanism could drive the shifts we observe. First, only a minority of voxels is expected to have ectopic receptive fields in the central representation; however our results show that at least 50% of voxels contribute to the displacements in the foveal representation. Secondly, silencing of ectopic neurons would ideally shrink the distribution of the neuronal population within a voxel, consequently, the position scatter should be away from the stimulus border i.e. parafoveal to foveal. Therefore, a more global neuronal mechanism might be inducing the observed changes in the pRF characteristics. Below, we outline how these variations might originate from extra-classical RF modulations and from attentional modulation.

### 4.3 Surround Suppression and Attention modulation

#### Surround suppression

fMRI-BOLD response amplitudes have been demonstrated to decrease with the introduction of a iso-oriented stimulus in the surround (Kastner et al., 2001; Nurminen et al., 2009; Williams et al., 2003; Zenger-Landolt and Heeger, 2003). In our case, the presence or absence of peripheral stimulation might in a way behave as the surround modulating the collective neural responses and shift the balance between excitatory (facilitation) and inhibitory (suppression) neuronal responses induced by the center-surround RF configurations (extra classical). Our results suggest that, for the masked conditions, the absence of a high-contrast iso-oriented surround mitigates the effects of surround suppression, resulting in the pRFs to be driven by much more excitatory neuronal activity.

#### Attention modulation

Previous studies reported response modulations in the visual cortex with voluntary attention and its influence in receptive field estimates (Desimone and Duncan, 1995; Kastner and Ungerleider, 2000; Klein et al., 2014; Puckett and DeYoe, 2015). In our study, the participants fixated on a cross in the center of the stimulus and performed an attention demanding task by reporting color changes of the fixation dot during all the conditions. In addition to this voluntary attention, masking the stimulus might induce an involuntary shift of attention towards the border of the stimulus (exogenous) and a shift in the balance between these two attentional modes might result in the pRF variations, we report here. Single cell studies in Macaque (Womelsdorf et al., 2008, 2006) showed that when attention is shifted from one location to another within the receptive field of the respective neuron, the RF centers of these neurons shift to the attended location. While these mechanisms, surround suppression and attentional field influence, are equally able to explain our data, the reported pRF variations might arise from an interaction of both mechanisms (Reynolds and Heeger, 2009) and other factors.

### 4.5 Limitations and future directions

At present we do not know which of the aforementioned mechanisms might be the most relevant to cause the observed variations in pRF measures, as our study was designed to identify the experimental effect, but not to pinpoint their exact origin. Consequently, further studies are needed to deduce them. With advances in pRF modeling approaches, a comparative study with other pRF modeling approaches might provide us with further insights into potential methodological/modeling biases. Forthcoming research should also investigate pRF data from patients and healthy controls with and without comparable experimental conditions (for e.g. artificial scotomas, visual acuity) to help us better understand and demonstrate the existence and impact of control biases in studies looking for potential cortical reorganization.

## 5. Conclusion

In summary, we demonstrated in healthy controls, the dependence of foveal pRF estimates on the spatial extent of the stimulation. We report enlargement and displacement of foveal pRFs towards the stimulus border when we reduced the size of the stimulus by masking and thereby restricting the peripheral stimulation. The shifts were more pronounced for the 4° than the 7° stimulus size and propagated across the primary (V1) and the extra striate (V2 & V3) visual cortex. Our results imply that, similar findings in patients with actual VF restrictions might also reflect normal cortical response behavior in a context of reduced peripheral stimulation. They therefore underscore that care must be taken to separate effects of stimulus properties, such as size, and of cortical reorganization in visual system pathologies. This emphasizes the importance of careful control measures in studies addressing neuronal plasticity.

## Data & code availability

The data and the code will be made available via a repository.

## Acknowledgements

This project was supported by European Union’s Horizon 2020 research and innovation programme under the Marie Sklodowska-Curie grant agreements No.675033 (EGRET plus), No 641805 (NextGenVis) and No. 661883 (EGRET cofund) to MBH & FWC. The funding organization did not have any role in the study design, collection, analysis and interpretation of the data, or publication of this research.

## CRediT author statement

Gokulraj Prabhakaran : Conceptualization, Methodology, Formal analysis, Investigation and data curation, Writing-original draft. Michael B. Hoffmann: Conceptualization, Methodology, Supervision, Writing-review and editing, Funding acquisition. Joana Carvalho: Software, Writing-review and editing. Azzurra Invernizzi: Software, Writing- review and editing. Martin Kanowski: Methodology, Writing- review and editing. Remco J. Renken: Software, Writing-review and editing. Frans W. Cornelisson: Software, Writing- review and editing, Funding acquisition.

