## Supplementary Figure 1 & 2 for "Foveal pRF properties in the visual cortex depend on the extent of stimulated visual field"

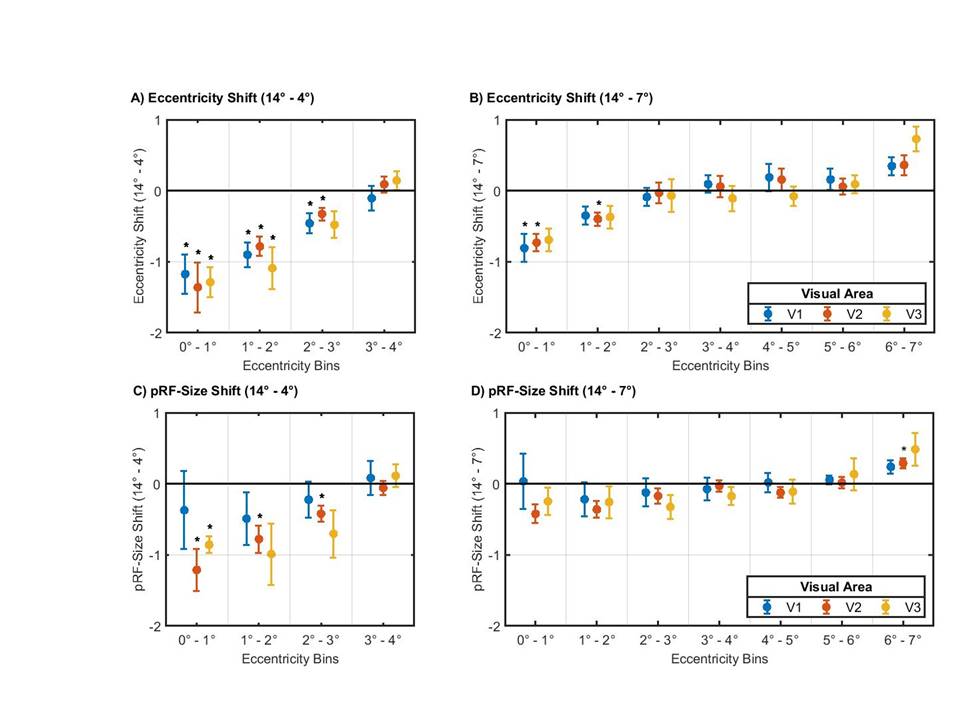

**Fig S1:** Stimulus size dependent differences in conventional pRF derived eccentricity and size; differences calculated as (**A)** Eccentricity_(14° - 4°)_ **(B)** Eccentricity_(14° - 7°)_ **(C)** pRF-size_(14° - 4°)_ **(D)** pRF-size_(14° - 7°)_. Grouping of voxels into eccentricity bins was performed as for Fig 3. Conventional pRF eccentricity and size estimates also exhibited shifts similar to those observed with the Bayesian approach in the foveal eccentricities (<3°) for the reduced stimulus size configurations. Visual areas, eccentricity bins and stimulus conditions with significant effects (p<0.05) after Holm-Bonferroni correction are indicated by “*”.

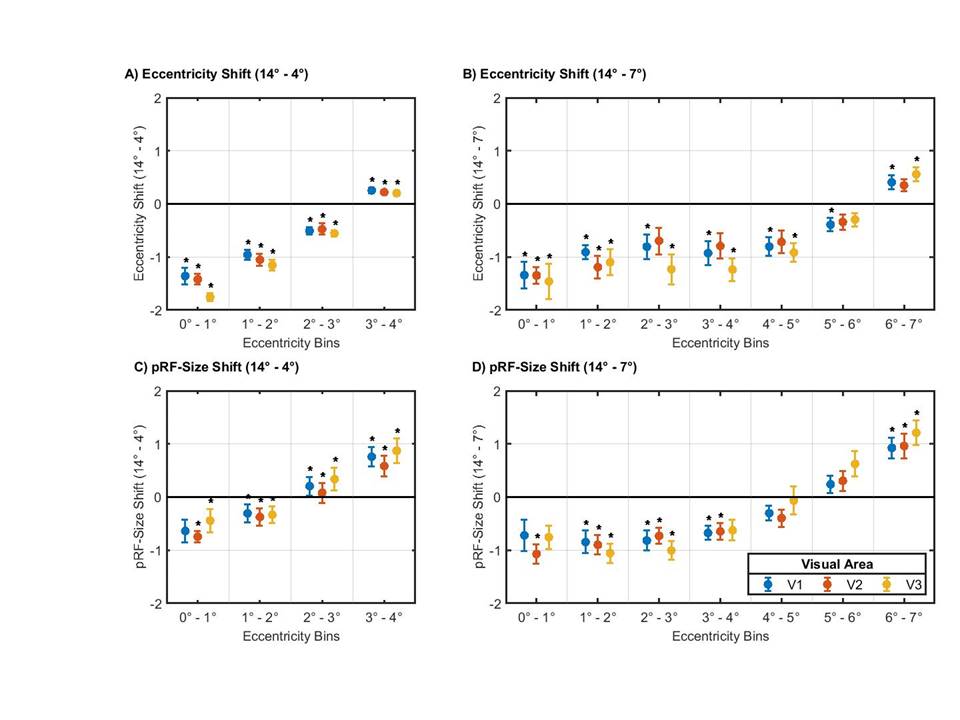

**Fig S2:** Stimulus size dependent differences in Bayesian pRF derived eccentricity and size when explicitly including the stimulus representation in the models for the masked stimulus configurations; differences calculated as (**A)** Eccentricity_(14° - 4°)_ **(B)** Eccentricity_(14° - 7°)_ **(C)** pRF-size_(14° - 4°)_ **(D)** pRF-size_(14° - 7°)_. Grouping of voxels into eccentricity bins was performed as for Fig 3. Significant shifts in eccentricity and pRF-Size were observed for the smaller stimulus conditions, similar to those observed when assuming an unmasked stimulus representation for all the models. However, the shifts were larger here and no longer restricted to the central eccentricities, in particular, for the 7° stimulus condition. Visual areas, eccentricity bins and stimulus conditions with significant effects (p<0.05) after Holm-Bonferroni correction are indicated by “*”.
